# MRI of OATP-expressing transplanted cells using clinical doses of gadolinium contrast agent

**DOI:** 10.1101/2023.08.07.552326

**Authors:** Tapas Bhattacharyya, Christiane L. Mallett, Jeremy M.-L. Hix, Erik M. Shapiro

## Abstract

Hepatic organic anion transporting polypeptides (OATPs) transport off-the-shelf, FDA-approved, hepatospecific Gd-based MRI contrast agents into cells that express the transporters enhancing signal on clinical T1-weighted MRI. Studies have used MRI to identify OATP-overexpressing tumors and metastases in mice following the delivery of Gd-EOB-DTPA at 27-67-fold higher than clinical doses. With safety and regulatory concern over Gd-based contrast agents, translation of this imaging paradigm to humans for regenerative medicine cell therapy will require substantially lower doses of agent.

Here we complemented static T1-weighted MRI and/or T1-mapping with dynamic contrast enhanced (DCE)-MRI and show that even low intracellular accumulation of contrast agent results in a sustained signal enhancement in OATP-overexpressing tumors while control, non-expressing tumors have rapid wash-in and wash-out dynamics which could be distinguished by performing area-under-the-curve (AUC) analyses.

## Introduction

Radiological monitoring will be a vital component of clinical regenerative medicine cell therapies and imaging reporter proteins will play key roles. In contrast to PET reporter proteins, MRI reporter proteins for imaging regenerative medicine cell therapies will be most useful when the high resolution and spatial discrimination capabilities of MRI are used for determining the precise anatomical locations of transplanted and long-term engrafted cells. This will be critical in regenerative medicine cell therapy, where clinicians need to know the precise delivered location of transplanted cells and track their persistence following *in vivo* expansion.

Hepatic organic anion transporting polypeptides (**OATPs**) have emerged as a promising, clinically viable MRI reporter protein concept^1-6^. Distinct amongst molecular imaging reporter protein paradigms, the translational potential lies in its simplicity – hepatic OATPs transport off-the-shelf, FDA-approved, hepatospecific Gd-based MRI contrast agents into cells that express the transporters^7-8^, the intracellular accumulation of which cells causes signal enhancement on clinical T1-weighted MRI^9^. Several groups have performed proof-of-principle studies, using MRI to identify OATP-overexpressing tumors in mice following the delivery of Gd-EOB-DTPA (Table 1). Indeed, following the delivery of Gd-EOB-DTPA, OATP-expressing tumors accumulate the contrast agent and become hyperintense on T1-weighted MRI, while control tumors do not. Excitingly, this has even been extended to locating distant metastases^1^.

**Table 1:** Studies reporting the use of OATPs for molecular imaging and the contrast agent (**CA**) dose used. **Note the clinical Gd-EOB-DTPA dose is 0.025 mmol/kg and all studies vastly exceed this**.

| Work | OATP type | Cell line | CA Dose | Image time | Ref |
| --- | --- | --- | --- | --- | --- |
| Patrick, 2014 | OATP1A1 | HCT-116 | 0.66 mmol/kg | 1 hour p.i. | 13 |
| Wu, 2018 | OATP1B3 | HT-1080 | 1.52 mmol/kg | 1 hour p.i. | 34 |
| Nystrom, 2019 | OATP1A1 | MDA-MB-231 | 0.1 mmol/kg | 5 hour p.i. | 32 |
| Wu, 2019 | OATP1B3 | PANC-1 | 1.52 mmol/kg | 1 hour p.i. | 33 |
| Kelly, 2021 | OATP1A1 | PC3 | 1.67 mmol/kg | 5 hour p.i. | 30 |
| Nystrom, 2023 | OATP1B3 | MDA-MB-231 | 1.3 mmol/kg | 5 hour p.i. | 31 |
| Wang, 2023 | OATP1B3 | Jurkat T cells | 1.5 mmol/kg | 5 hour p.i. | 35 |

Yet, these early proof-of-principle successes come with an important caveat. The standard clinical dose for Gd-EOB-DTPA is 0.025 mmol/kg ^9^, while most OATP reporter protein studies have used 27-to 67-fold higher doses^2-4, 6^ (Table 1). Also, because of the high dose of contrast agent delivered, studies have had to wait 1-5 hours post injection to perform MRI, to allow the substantial non-accumulated agent to be cleared from the blood. With recent safety and regulatory concern over Gd-based contrast agents ^10^, translation of this imaging paradigm to humans will require substantially lower doses of contrast agent than what has been reported.

In this work, we evaluated two strategies for using MRI to specifically identify OATP-overexpressing tumors using the standard clinical dose of MRI contrast agent. The first strategy was to use a rat OATP1B2 as the reporter protein, as rat OATP1B2 efficiently transports both Gd-BOPTA and Gd-EOB-DTPA, while the human liver poorly transports Gd-BOPTA^9^. The rationale for this was that by using a Gd-BOPTA-transporting OATP, we could inject a lower dose of Gd-BOPTA as hepatic background uptake would be low, while tumor uptake would remain high. These experiments were performed in a mouse model expressing human hepatic OATPs rather than rodent OATPs, mimicking an experiment performed in a human^11^. The second strategy was to complement static T1-weighted MRI and/or T1-mapping with dynamic contrast enhanced (DCE)-MRI and use the dynamic data to distinguish OATP-overexpressing tumors from control non-expressing tumors via differential contrast agent dynamics. Based on previous studies showing prolonged retention of contrast agent within OATP-overexpressing tumors^3, 6^, we hypothesized that even low intracellular accumulation of contrast agent would result in a sustained elevation of signal intensity in OATP-overexpressing tumors while control, non-expressing tumors would have a rapid wash-in and wash-out dynamic which could be distinguished by simply performing area-under-the-curve (AUC) analyses^12^. Ultimately, we show that AUC analysis of the DCE-MRI curves can robustly distinguish OATP-overexpressing tumors from non-expressing tumors, and importantly, background tissue, using standard clinical dose of MRI contrast agent. We did not find that using rat OATP1B2, coupled with Gd-BOPTA provided an advantage over using Gd-EOB-DTPA, at least in this animal model.

## Material & Method

### Stable cell lines

Stable cell line constitutively expressing OATP1B2 were created by lentiviral transduction follow by the selection and enrichment by antibiotic selection. Lentiviral vector carrying codon optimized rat OATP1B2 under the control of EF1A promoter and blasticidine (Bsd) resistance gene under the control of mPGK promoter was purchased from Vector Builder. Human OATP1B3 and pig OATP1B4 were also similarly designed and purchased. MyC-Cap cells (mouse prostate cancer cells) and HEK-293 (human embryonic kidney cell lines) were purchased from ATCC. Both the cell lines were infected separately with 10 MOU viral particles followed by antibiotic selection at sublethal concentration. Standard kill kinetic assay was performed for both the cell lines against blasticidine to determine the concentration of antibiotic for selection. MyC-Cap and HEK-293 cells were selected at a concentration of 5 μg/ml and 10 μg/ml respectively.

### In-vitro MRI

Cells were incubated in culture media with either Gd-EOB-DTPA or Gd-BOPTA (2.5 mM, 1 hour). Media was then removed, cells were washed with PBS, and then trypsinized to detach from the plate. Cell pellets were washed twice more with PBS before MRI (Bruker 7.0T, 70/30 BioSpec). T1 maps of cell pellets were measured using standard sequences. Cells with lowest T1 values (highest contrast agent uptake) progressed to in vivo studies.

### Animal Model

Tumors were developed by subcutaneous injection of 2×10^6^ cells in 100ul PBS with 100ul Matrigel on the right (OATP expressing cells) and left flanks (wild-type cells) of mice. Humanized OATP knock-in mice^11^ or wild-type FVB mice were used to evaluate the background effects of human and rodent transporters in the liver. Mice underwent MRI two weeks post-inoculation, when tumors had formed.

### In-vivo MRI

Mice underwent MRI on a Bruker 70/30 BioSpec using a 40 mm volume coil. Contrast agent doses for Gd-EOB-DTPA were 0.025 mmol/kg (clinical dose) or 0.25 mmol/kg (10x clinical dose), and for Gd-BOPTA were 0.025 mmol/kg (0.5 clinical dose) or 0.25 mmol/kg (5x clinical dose). Before tail-vein IV contrast agent administration, T2-weighted RARE (TR/TE 3000/40 ms) and T1-weighted RARE (TR/TE 750/6.5 ms) images were acquired. A dynamic T1-weighted FLASH sequence (TR/TE 46/2.5 ms) was acquired during injection and continuing for 1 hour with 30 s temporal resolution and then a T1-weighted RARE image was acquired 1 hour post contrast administration. Image resolution was 200×200 um and slice thickness was 1mm for all sequences.

## Results

Rat OATP1B2 overexpressing MyC-Cap cells had the lowest T1 values (Figure 1), so these cells progressed to in vivo experiments. Figure 2 shows MRI from a humanized OATP-knock-in mouse harboring a rat OATP1B2-overexpressing tumor and a wild-type tumor. This mouse received 10X clinical dose of Gd-EOB-DTPA. Substantial signal enhancement in the OATP-overexpressing tumor is observed. Figure 2 shows MRI from a wild-type FVB mouse harboring a rat OATP1B2-overexpressing tumor and a wild-type tumor. This mouse received half clinical dose of Gd-BOPTA. In this example, hyperintense MRI signal is less obvious. AUC maps were abot to discriminate between wild-type and OATP-1B2 expressing tumors after low dose of Gd-BOPTA. In Figure 3 the DCE-MRI time course clearly demonstrates the retained low hyperintensity on the OATP-overexpressing tumor versus wash out from the wild-type tumor. Signal enhancement is patchy because of the way these tumors grow. Figure 4 demonstrates that acquisition times of 30-45 minutes are needed to discriminate between wild type tumors and OATP1B2-expressing tumors. More careful ROI analysis to include only tumors will likely reduce this time further.

**Figure 1:**
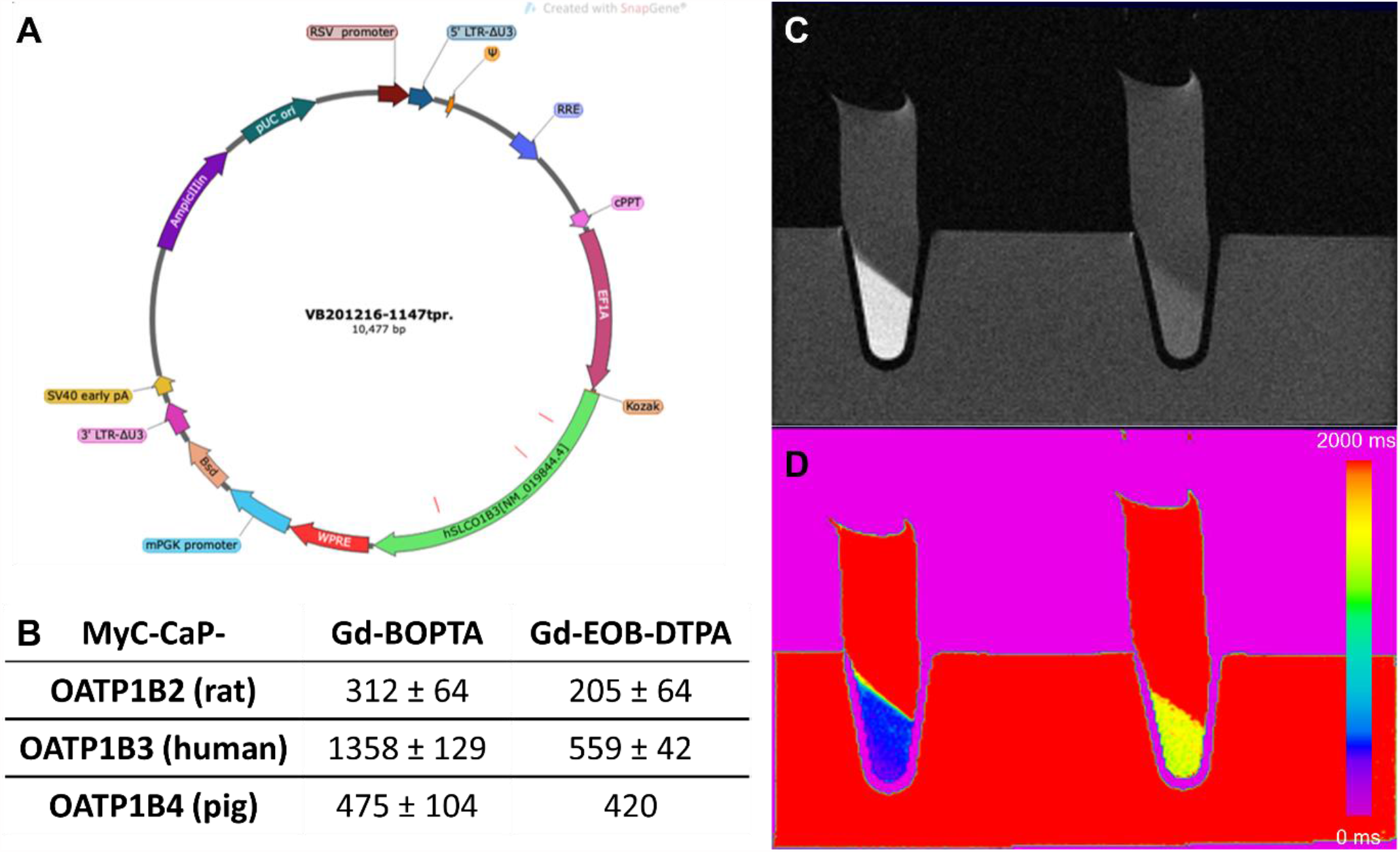
In vitro creation and validation of MyC-CaP-OATP cells. A: Schematic of lentiviral vector. B: Comparison of T_1_ for 3 different OATPs. OATP1B2 had the lowest T1 so was used for further studies. C: Sample T_1_ weighted image of a transduced (left) and a wild-type (right) cell pellet after incubation with Gd-BOPTA. D: T1 map for the same pellets as (D). The T_1_ values are 400 ms and 1525 ms.

**Figure 2:**
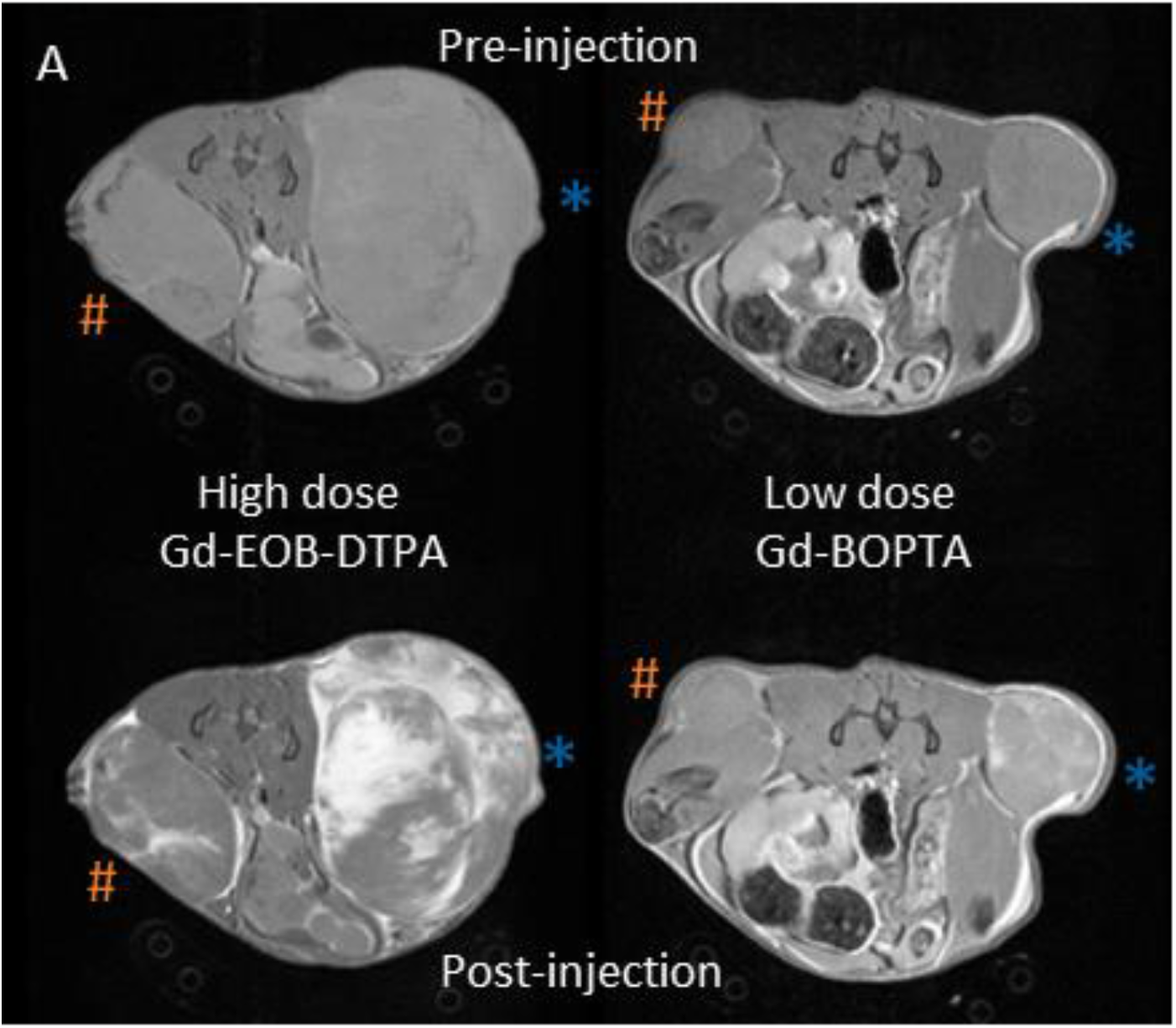
Transduced cells are detectable after low-dose GBCA injection using area under the curve mapping from dynamic MRI. A: On T1 weighted images, transduced tumor is very bright 1h after injection of high dose Gd-EOB-DTPA but difference between tumors is slight after low dose Gd-BOPTA.

**Figure 3:**
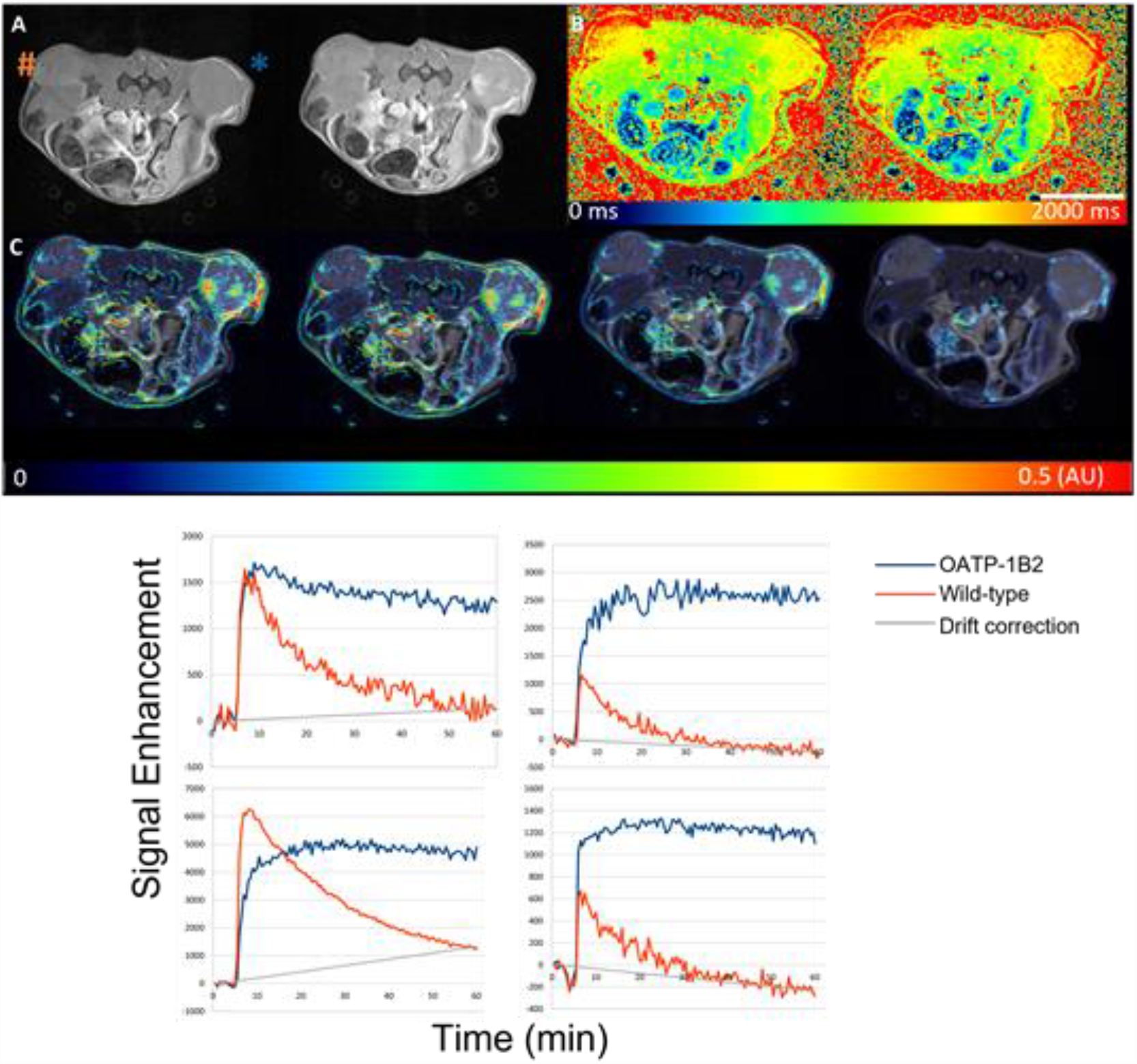
Top) AUC maps are the best way to discriminate between wild-type and OATP-1B2 expressing tumors after low dose of Gd-BOPTA. A: T1 weighted images before and after injection. B: T1 maps, before and after injection. C: AUC maps at 0-60, 0-45, 0-30 and 0-15 mins. Tumors can be discriminated at 30 mins. Scale bar is 1 cm. Orange hash is wild-type tumor, blue asterisk is OATP-1B2 tumor. Bottom) **Sample signal enhancement curves for 1x Gd-EOB-DTPA and Gd-BOPTA in several mice**. AUC was calculated as the area between the curves and the grey line, which was used to compensate for signal drift due to heating during MRI. This need for this can be removed by using drift compensation in future acquisitions.

**Figure 4:**
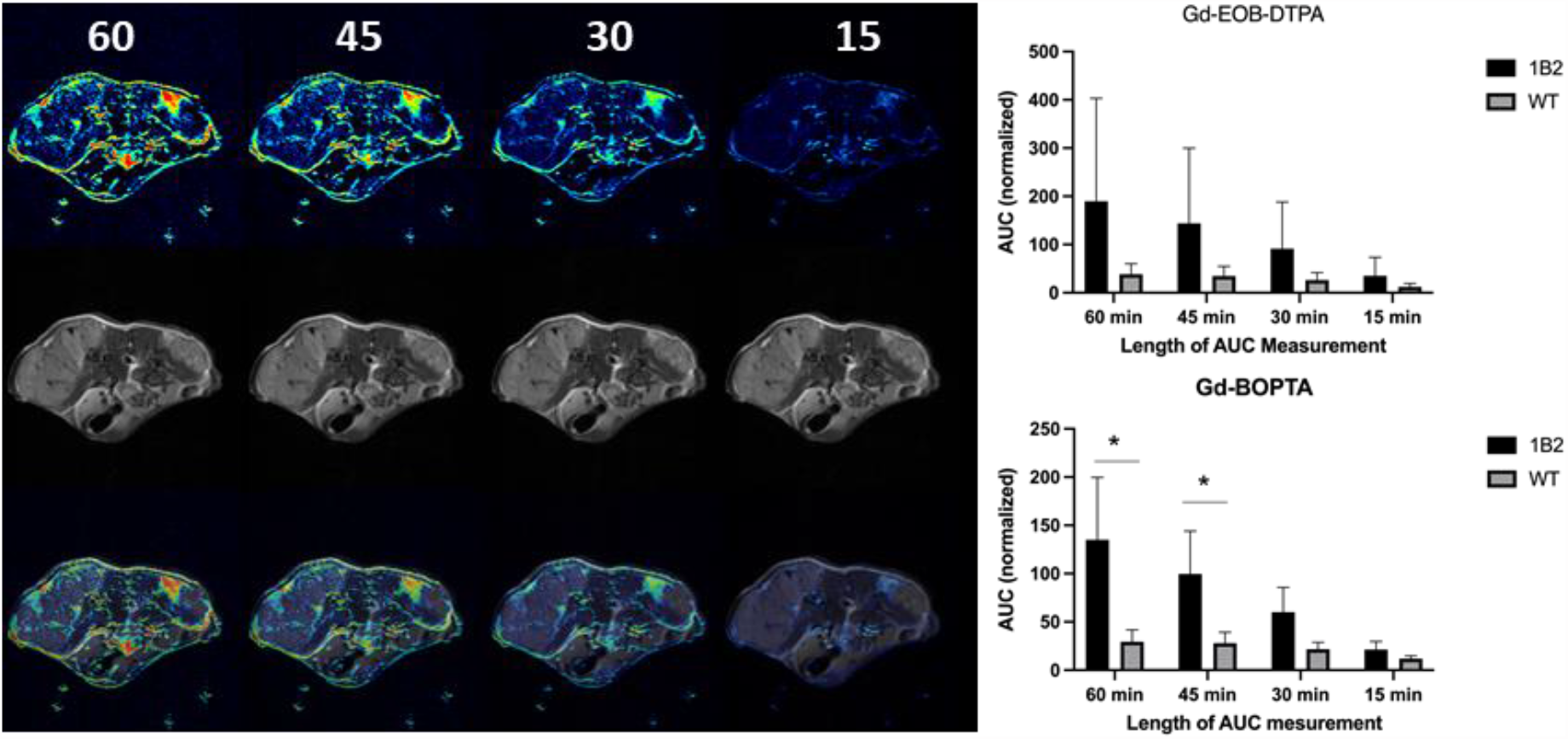
Left) AUC maps of mouse injected with 1X Gd-BOPTA at different time points. Right) AUC was different between tumor types after Gd-BOPTA injection at 60 and 45 minutes. For Gd-EOB-DTPA, there was a trend toward significance (p<0.1). The AUC is normalized to the peak enhancement of the wild-type tumor.

Rat OATP1B2 overexpressing cells grew similarly in humanized OATP mouse models as in syngeneic FVB mice, suggesting that rat OATP1B2 has low immunogenicity in genetically disparate scenarios.

## Discussion and Conclusion

A major challenge in cell/gene therapy is to monitor therapeutic/engineered cells through non-invasive imaging. MRI reporter genes and clinically approved contrast agents can potentially solve this dilemma, however, agents need to be delivered at clinical doses or lower for clinical translation. We first screened species-specific hepatic OATPs in transduced cell lines and determined that rat OATP1B2 was the most effective OATP for transporting both Gd-EOB-DTPA and Gd-BOPTA. Rat OATP1B2 is 63 percent (456 residues) identical to human OATP1B3 with 161 additional similar residues, yet we acknowledge the potential challenge of using a rodent protein for human molecular imaging. Screening of OATP-expressing cell lines for maximal accumulation of agent via in vitro T1 mapping enabled rational choice of cell line for generating OATP-expressing tumors that could be detected in vivo in mice using clinical dose of agent. The combination of T1-weighted MRI and DCE-MRI facilitates the differentiation of OATP-expressing tumors where T1-weighting alone may not be sufficient. Thus, OATP-overexpressing cell transplants can be detected in vivo using standard clinical doses and pixelwise mapping of AUC can be used to identify cell transplants with persistent uptake of hepatospecific contrast agents at clinical doses.

